# m^1^A58 acts as a conformational checkpoint coupling human initiator tRNA maturation to translation initiation

**DOI:** 10.64898/2026.08.28.747798

**Authors:** Hao Li, Xing-Yi Wu, You-Jia Zhou, Marcel-Joseph Yared, Chen-Xi Wang, Pei-Yu Tian, Qing-Yun Liu, Zhong-Gui Bao, Pierre Barraud, Ru-Juan Liu

## Abstract

tRNAs are characterized by extensive chemical modifications that influence tRNA fate. N^1^-methyladenosine at position 58 (m^1^A58) is a widespread core tRNA modification linked to physiological and pathological processes. However, how m^1^A58 coordinate tRNA folding and processing to ensure translational efficiency in mammalian cells remains largely unknown. Using acute dTAG-mediated degradation and CRISPR-Cas9 knockout, we identified initiator methionine tRNA (tRNA^iMet^) as selectively vulnerable to m^1^A58 loss, lacking the isodecoder buffering observed for most other tRNA isoacceptors. NMR analysis of the tRNA^iMet^ showed that m^1^A58 stabilizes D/T-loop interactions, consistent with a maturation-competent conformation. *In vitro* processing assays further demonstrated that m^1^A58 promotes RNase P-mediated 5’-leader removal and RNase Z-mediated 3’-trailer cleavage, while La/SSB protects accumulated precursors. Disrupting this checkpoint impaired the assembly of the eIF2-containing 43S pre-initiation complex and global protein synthesis, which was substantially rescued by adding m^1^A58-modified tRNA^iMet^. Acute TRMT6 degradation elicited temporally coordinated gene-expression responses involving proteostasis, transport and signaling. Together, these findings establish m^1^A58 as a conformational checkpoint coupling human initiator-tRNA maturation to translation initiation and stress responses.

## Introduction

tRNAs carry extensive chemical modifications that shape their processing, stability and translational function (Pan, 2018; Phizicky & Hopper, 2023; Schimmel, 2017; Suzuki, 2021; Zhu *et al*, 2026). Dysregulated tRNA modifications have been linked to tumor progression, neurological disease, immune regulation, metabolic homeostasis and development (Chujo & Tomizawa, 2025; Delaunay *et al*, 2023; Delaunay *et al*, 2022; Hughes *et al*, 2024; Kirchner & Ignatova, 2014; Motorin & Helm, 2010; Orellana *et al*, 2022; Suzuki, 2021; Torres *et al*, 2014; Zhang & Lu, 2024). Among these modifications, N^1^-methyladenosine at position 58 (m^1^A58) is one of the most widespread and evolutionarily conserved tRNA core modifications, found across all domains of life (Björk *et al*, 1987; Boccaletto *et al*, 2021; He *et al,* 2024; Saikia *et al*, 2010). In eukaryotic cytosolic tRNAs, m^1^A58 is generally installed by the conserved Trm6/Trm61 methyltransferase complex, known as TRMT6/TRMT61A in mammals (Anderson *et al*, 1998; Anderson *et al*, 2000; Li *et al*, 2017; Ozanick *et al*, 2005; Safra *et al*, 2017; Wang *et al*, 2016). m^1^A58 is a reversible tRNA mark that dynamically controls tRNA stability and translation efficiency through its deposition and removal (Alexandrov *et al*, 2006; Chernyakov *et al*, 2008; Dewe *et al*, 2012; Kadaba *et al*, 2004; Liu *et al*, 2016; Maraia *et al*, 2022; Nguyen *et al*, 2025; Roundtree *et al*, 2017; Zhang & Jia, 2018). Some studies further indicate that TRMT6/TRMT61A-mediated tRNA m^1^A58 contributes to tumorigenesis and cancer progression across multiple cancer types (Macari *et al*, 2015; Tao *et al*, 2025; Wang *et al*, 2021) and is also required for efficient T cell activation (Liu *et al*, 2022), stem cell maintenance (Zuo *et al*, 2024) and and HIV-1 reverse transcription and replication (Renda *et al,* 2001). Despite the importance of m^1^A58 in both physiological and pathological contexts, the molecular mechanisms by which it controls tRNA fate in mammalian cells remain poorly understood.

Pol III-transcribed pre-tRNAs undergo a multistep maturation process involving RNase P-mediated 5’-leader removal (Elder *et al*, 2024; Hopper, 2013; Phizicky & Hopper; Stanley *et al*, 2026; Wu *et al*, 2018), RNase Z/ELAC2-mediated 3’-trailer cleavage (Bhatta *et al*, 2025; Brzezniak *et al*, 2014; de la Sierra-Gallay *et al*, 2005; Elder *et al*, 2024; Hopper, 2013; Stanley *et al*, 2026), 3’-CCA addition (Phizicky & Hopper, 2023; Stanley *et al*, 2026), and extensive nucleotide modification (Peschek & Tuorto, 2025; Schultz & Kothe, 2024; Stanley *et al*, 2026). In *Saccharomyces cerevisiae*, m^1^A58 is critical for tRNA^iMet^ biogenesis (Anderson *et al*, 1998; Basavappa & Sigler, 1991; Calvo *et al*, 1999; Kadaba *et al*, 2004; Ozanick *et al*, 2009; Schneider *et al*, 2007; Tasak & Phizicky, 2022; Wang *et al*, 2008; Yared *et al*, 2023). Loss of m^1^A58 in yeast triggers degradation of pre-tRNA^iMet^ through the nuclear surveillance pathway and the rapid tRNA decay (RTD) pathway (Kadaba *et al*, 2004; Kramer & Hopper, 2013; Li *et al*, 2016; Maraia *et al*, 2022; Ozanick *et al*, 2009; Phizicky & Hopper, 2023; Schneider *et al*, 2007; Tasak *et al*, 2022; Wang *et al*, 2008). The nuclear surveillance pathway involves Rex1p and the TRAMP-exosome machinery, which act on hypomodified pre-tRNA^iMet^ (Kadaba *et al*, 2006; Ozanick *et al*, 2009; Schneider *et al*, 2007; Wang *et al*, 2008). Our recent work further showed that loss of m^1^A58 induces an aberrant conformation of yeast tRNA^iMet^, which may facilitate its recognition by RNA quality-control pathways (Yared *et al*, 2023). Notably, human tRNA^iMet^ is fully modified at m^1^A58 across all tested cell lines, consistent with its role in tRNA^iMet^ stability (Saikia *et al,* 2010). Moreover, m^1^A58 has also been detected on pre-tRNAs in human cells (Cozen *et al,* 2015), indicating that this modification can be installed before completion of tRNA maturation. However, how m^1^A58 regulates tRNA biogenesis in mammalian cells remains poorly understood.

tRNA maturation is governed by both enzymatic processing and higher-order folding, with the conserved elbow stabilizing the canonical L-shaped fold (Basavappa & Sigler, 1991; Peschek & Tuorto, 2025; Schultz & Kothe, 2024; Robertus *et al,* 1974; Suddath *et al,* 1974; Zhang & Ferré-D’Amaré, 2016). Structural studies support this architecture-based view: human nuclear RNase P engages the acceptor-T-arm domain and elbow region (Elder *et al*, 2024; Wu *et al*, 2018), whereas RNase Z contacts the T-arm/elbow region through its flexible arm to promote accurate 3’ end processing (Bhatta *et al*, 2025; Brzezniak *et al*, 2014; Elder *et al*, 2024; Xue *et al*, 2024). m^1^A58 is positioned within the T-loop/elbow structural core, placing it at a critical node linking tRNA modification, folding and processing (Yared *et al*, 2023). Our work showed that, in *in vitro*-transcribed yeast tRNA^iMet^, introduction of m^1^A58 alone markedly increases structural homogeneity and promotes proper elbow assembly (Yared *et al*, 2023), suggesting that m^1^A58 may function as a folding-coupled maturation checkpoint. However, it remains unclear whether loss of m^1^A58 induces analogous conformational perturbations in mammalian tRNAs and whether these perturbations further affect pre-tRNA end processing and the subsequent fate of processed tRNA.

Here, we combine acute TRMT6 depletion, genetic perturbation, quantitative tRNA profiling, NMR and biochemical assays to define how m^1^A58 controls human tRNA maturation. We identify human tRNA^iMet^ as a modification-sensitive substrate in which m^1^A58 establishes a processing-competent conformation required for efficient RNase P- and RNase Z-mediated end maturation. Loss of this state impairs end processing and promotes accumulation of La/SSB-bound precursors, thereby reducing the availability of functional mature tRNA. Notably, most tRNA isoacceptors are buffered against m^1^A58 loss through their abundant and stable isodecoder members. tRNA^iMet^, however, lacks such resilience because its isodecoders are highly abundant but intrinsically short-lived and rapidly turned over. Time-resolved transcriptomic analyses following acute TRMT6 degradation further reveal temporally ordered gene-expression remodeling, involving dynamic regulation of proteostasis, transport, and signaling pathways. Together, our study defines m^1^A58 as a conformational maturation checkpoint that links tRNA folding and processing to translation initiation and cellular stress responses in human cells.

## Results

### Establishment of an acute TRMT6/TRMT61A degradation system and stable *TRMT6*-deficient cell models

Previous studies showed that deletion of Trmt6 in mice causes severe hematopoietic failure and eventual lethality (Zuo *et al*, 2024). This *in vivo* phenotype highlights the importance of TRMT6 in mammalian cells. To enable rapid and inducible depletion of TRMT6 or TRMT61A, we generated endogenous dTAG-*TRMT6* and *TRMT61A*-dTAG systems in HEK293T cells (Fig. 1A) (Nabet *et al*, 2018). Specifically, a dTAG degron (comprising the FKBP12^F36V^ tag) was introduced into the endogenous *TRMT6* or *TRMT61A* locus together with a puromycin resistance cassette, allowing selection of stable cells. In this system, addition of the dTAG ligand (dTAG-13) recruits endogenous TRMT6 or TRMT61A to the ubiquitin-proteasome degradation pathway, thereby enabling temporally controlled acute protein depletion (Fig. 1A) (Nabet *et al*, 2018). To validate the degradation systems, dTAG-TRMT6 and TRMT61A-dTAG cells were treated with dTAG-13 for 48 h and subjected to Western blot analysis. In both systems, the corresponding target protein was efficiently depleted (Fig. 1B).

**Figure 1:**
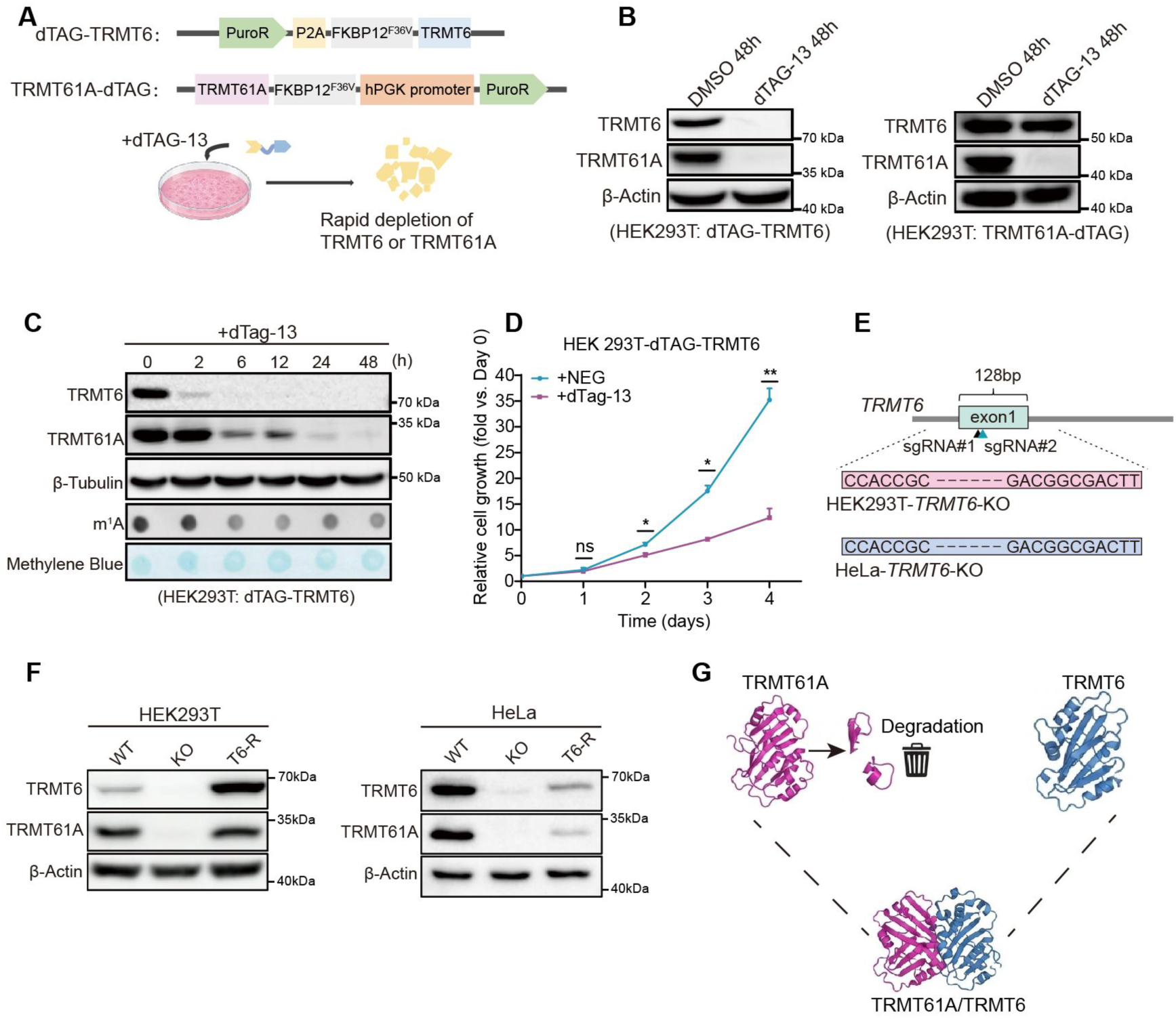
Generation of TRMT6/TRMT61A degron and TRMT6 knockout cell models. **(A)** Schematic of the endogenous dTAG-TRMT6 and TRMT61A-dTAG constructs generated in HEK293T cells. FKBP12^F36V^ was fused to the N terminus of TRMT6 or the C terminus of TRMT61A. Puromycin-resistance cassettes were included for selection. Addition of dTAG-13 induces rapid proteasomal degradation of the tagged protein. **(B)** Western blot analysis of TRMT6 and TRMT61A in HEK293T dTAG-TRMT6 cells (left) and TRMT61A-dTAG cells (right) treated with DMSO or dTAG-13 for 48 h. β-Actin served as a loading control. **(C)** Time-course analysis of TRMT6 degradation. HEK293T dTAG-TRMT6 cells were treated with dTAG-13 and collected at the indicated time points (0-48 h). TRMT6 and TRMT61A protein levels were examined by Western blotting, with β-Tubulin as the loading control. Total RNA m^1^A levels were assessed by dot blot, and methylene blue staining was used to verify RNA loading. **(D)** ATP-based measurement of relative cell growth in HEK293T dTAG-*TRMT6* cells treated with dTAG-13 or the negative control (+NEG) for the indicated number of days. Values are presented relative to day 0. Statistical significance is indicated in the graph: ns, not significant; *P < 0.05; **P < 0.01. **(E)** CRISPR–Cas9 strategy used to generate TRMT6-knockout HEK293T and HeLa cells. Two sgRNAs targeting exon 1 of TRMT6 were used to delete the indicated 128-bp genomic region. **(F)** Western blot validation of *TRMT6*-knockout cells. TRMT6 and TRMT61A protein levels were examined in wild-type (WT), *TRMT6*-knockout (KO), and TRMT6-reconstituted (T6-R) HEK293T and HeLa cells. β-Actin served as a loading control. **(G)** Proposed model illustrating the asymmetric stability relationship within the TRMT6/TRMT61A complex.

To examine the time course of dTAG-13-mediated TRMT6 depletion, dTAG-TRMT6 cells were treated with dTAG-13 and harvested at the indicated time points for Western blot analysis. TRMT6 protein levels were markedly reduced after 2 h of dTAG-13 treatment and became nearly undetectable by 6 h (Fig. 1C). Consistent with TRMT6 depletion, total RNA m^1^A levels, as measured by dot blot, began to decrease at 6 h (Fig. 1C). These results indicate that this system efficiently depletes TRMT6 within a short time window and rapidly reduces TRMT6-associated RNA methylation. Notably, although TRMT6 was rapidly depleted within 6 h, cells remained viable after 72 h of treatment or longer (Fig. 1D). This result indicates that acute TRMT6 depletion does not immediately cause cell death in HEK293T cells. This contrasts with the severe phenotype observed after prolonged TRMT6 loss *in vivo* and may reflect the different consequences of acute versus chronic TRMT6 deficiency.

To assess long-term tolerance to TRMT6 loss and distinguish acute depletion effects from chronic adaptive phenotypes, we next generated *TRMT6* knockout cells in both HEK293T and HeLa cells (Fig. 1E). Candidate clones obtained by genome editing were validated by Western blotting (Fig. 1F), which confirmed loss of detectable TRMT6 protein and successful establishment of *TRMT6* KO cells in both cell lines. The ability of TRMT6 KO cells to be established and expanded further supports that TRMT6 loss is not immediately lethal in these cultured cell models, although it may induce remodeling of cellular state and gene-expression programs.

During validation of the TRMT6-dTAG and *TRMT6*-KO models, we examined TRMT61A, the catalytic partner of TRMT6, and found that its protein levels decreased in parallel with TRMT6 following dTAG-induced degradation and were also markedly reduced in *TRMT6*-KO cells (Fig. 1B, C, F). Re-expression of TRMT6 restored TRMT61A protein levels (Fig. 1F), whereas acute TRMT61A degradation had little effect on TRMT6 abundance (Fig. 1B), revealing an asymmetric dependence of TRMT61A protein stability on TRMT6 (Fig. 1G).

Together, these data validate our acute and stable cell models for TRMT6 and TRMT61A depletion, and further reveal that TRMT61A protein stability depends on TRMT6.

### Anticodon-defined tRNA pool remains largely stable against m^1^A58 loss except for tRNA^iMet^

m^1^A58 is widely present in human cytoplasmic tRNAs and has important regulatory functions (Saikia *et al*, 2010). However, how m^1^A58 loss affects the composition and homeostasis of the human tRNA pool remains unclear. To determine how m^1^A58 loss affects the human tRNA pool, we performed tRNA sequencing (tRNA-seq) in both *TRMT6*-dTAG and *TRMT6*-knockout cells (Fig. 2A). In the dTAG system, cells were collected at 0, 2, 4, 24, and 72 h after dTAG-13 treatment. First, we estimated m^1^A58 levels based on the mismatch frequency at position 58 generated during reverse transcription (RT). The overall m^1^A58-associated mismatch signal began to decrease by 4 h, declined progressively thereafter, and was nearly undetectable by 72 h (Fig. EV1A-F). Next, we quantified tRNA abundance at the anticodon level by grouping tRNA species sharing the same anticodon and calculated Pearson correlation coefficients between the 0 h control and each subsequent time point (2, 4, 24, and 72 h), yielding values of 0.995, 0.967, 0.986, and 0.958, respectively (Fig. 2B-E). These results suggest that, despite the progressive loss of m^1^A58, the overall tRNA abundance profile at the anticodon level remained largely stable, with only modest changes at later time points (Fig. 2B-E). Similarly, anticodon-level tRNA abundance was also strongly correlated between WT and stable *TRMT6*-KO cells (Pearson r=0.930; Fig. 2F). This correlation was slightly lower than those observed in the dTAG system, suggesting a greater degree of tRNA-pool remodeling in stable *TRMT6*-KO cells. Together, these results showed that the overall tRNA abundance profile at the anticodon level was broadly buffered against both acute m^1^A58 loss and stable *TRMT6* knockout, with only modest perturbations to the overall tRNA pool.

**Figure 2:**
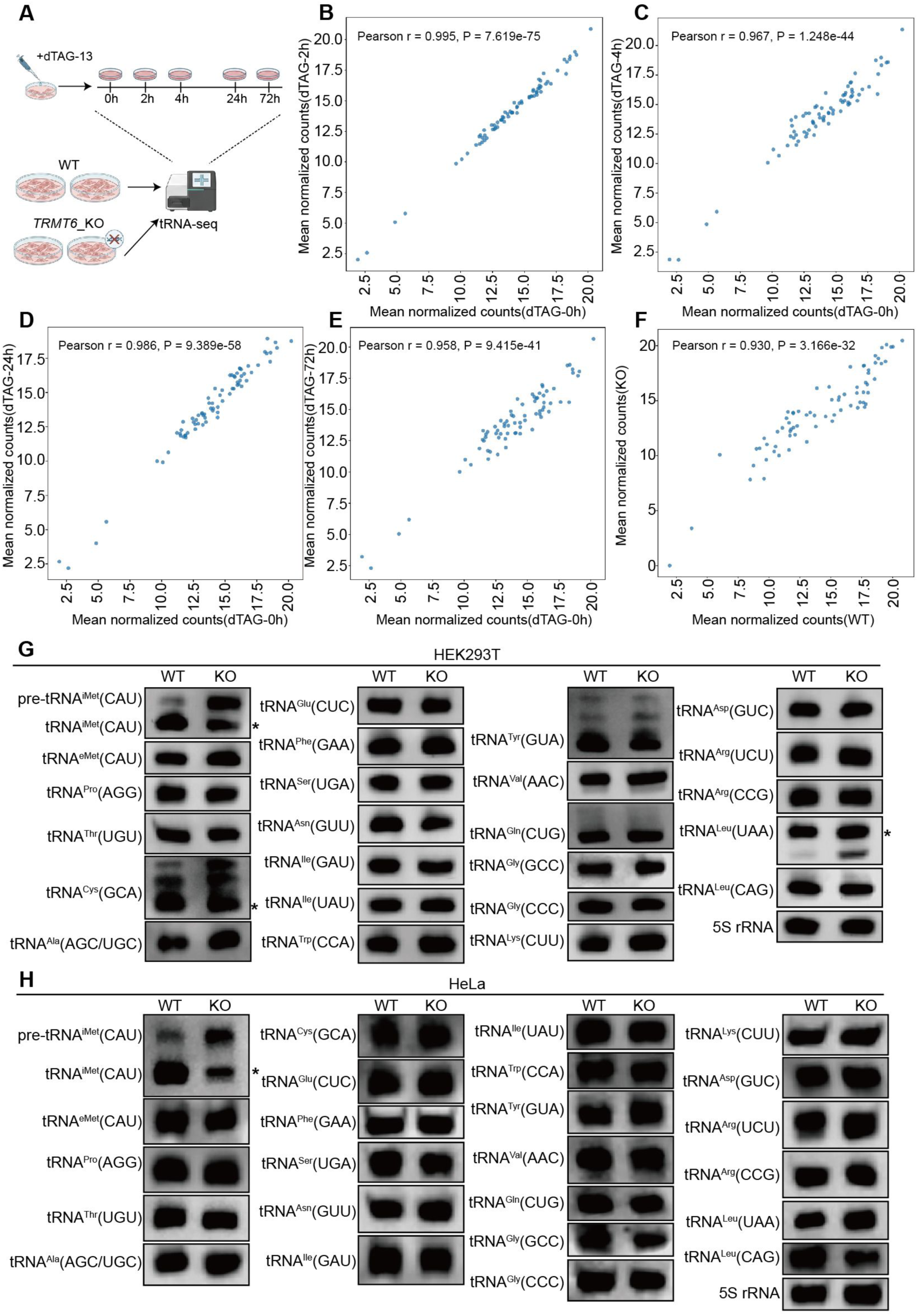
Time-resolved tRNA profiling reveals buffering at the anticodon-pool level following m^1^A58 loss. **(A)** Schematic of the experimental design. Degron-engineered cells were treated with dTAG-13 and collected at 0, 2, 4, 24, and 72 h for tRNA sequencing (tRNA-seq). In a separate experiment, wild-type (WT) and *TRMT6*-knockout (KO) cells were subjected to the same tRNA-seq workflow. **(B-E)** Pairwise correlation analyses of anticodon-pool abundance between degron cells at 0 h and those collected at 2 h (B), 4 h (C), 24 h (D), or 72 h (E) after dTAG-13 treatment. Each point represents one anticodon-defined tRNA pool, plotted using its mean normalized tRNA-seq count. **(F)** Correlation of anticodon-pool abundance between WT and *TRMT6*-KO cells. Pearson correlation coefficients (r) and corresponding P values are shown in each panel. **(G, H)** Northern blot validation of the indicated precursor and mature tRNA species in WT and *TRMT6*-KO HEK293T **(G)** and HeLa **(H)** cells. 5S rRNA served as a loading control. Asterisks (*) indicate the expected migration positions of mature tRNAs based on their predicted sizes.

To validate the tRNA-seq results, we performed Northern blotting in WT and *TRMT6*-KO HEK293T and HeLa cells. We examined 24 representative tRNA species or families that could be reliably detected (Fig. 2G, H). Most mature tRNAs showed no detectable changes in *TRMT6*-KO cells. These findings were consistent with the overall stability observed by tRNA-seq. However, tRNA^iMet^(CAU), which serves as a core component of the 43S pre-initiation complex, was a notable exception. Its mature form was markedly reduced in *TRMT6*-KO cells, whereas its precursor accumulated in both HEK293T and HeLa cells (Fig. 2F-H). This reduction in mature tRNA^iMet^ was also consistent with the tRNA-seq results (Fig. EV2A). In contrast, elongator tRNA^eMet^(CAU) remained largely unchanged (Fig. 2F-H). Thus, not all tRNAs were sensitive to m^1^A58 loss; instead, the tRNA^iMet^ exhibited the most prominent and selective response. Consistent with this, northern blotting in the dTAG system confirmed that mature tRNA^iMet^ began to decline within 6 h of dTAG-13 treatment, whereas its precursor accumulated as early as 2 h, further supporting a rapid maturation defect upon acute m^1^A58 loss (Fig. EV2B). The accumulation of its precursor and the reduction of its mature form suggest that m^1^A58 is required for the efficient maturation and/or stability of initiator tRNA^iMet^.

Together, these results establish that the human tRNA pool is globally stable against m^1^A58 loss at the anticodon level, with the notable exception of tRNA^iMet^. Its exquisite sensitivity likely reflects unique structural constraints that render its maturation critically dependent on this modification.

### Lack of isodecoder buffering renders tRNA^iMet^ vulnerable to m^1^A58 loss

Each anticodon-defined tRNA family comprises multiple isodecoders that share the same anticodon but differ at other positions in their sequences. For example, the tRNA^iMet^(CAT) family consists of three isodecoders (Fig. 3A). To investigate why tRNA^iMet^ is particularly sensitive to m^1^A58 loss, we compared tRNA abundance between *TRMT6*-knockout and wild-type cells at the isodecoder level (Fig. 3B). tRNA-seq revealed that individual isodecoders responded differently to TRMT6 loss (Fig. 3B). Within the tRNA^iMet^(CAT) family, the two readily detectable isodecoders, tRNA^iMet^(CAT)-1 and tRNA^iMet^(CAT)-2, were significantly reduced in *TRMT6*-knockout cells. Because tRNA^iMet^(CAT)-3 was expressed at an extremely low level, it was excluded from subsequent quantitative analyses (Fig. 3B). By contrast, in most other tRNA families, only one or a small subset of the constituent isodecoders showed significant changes in abundance, whereas the remaining members showed no significant changes (Fig 3B; Appendix Table S1). For example, among the 16 tRNA^Asn^(GTT) isodecoders, only tRNA^Asn^(GTT)-24 was significantly reduced (Fig. 3B; Appendix Table S1). Likewise, among the eight tRNA^Tyr^(GTA) isodecoders, only tRNA^Tyr^(GTA)-1 was significantly increased (Fig. 3B; Appendix Table S1). Thus, TRMT6 loss generally affected specific isodecoders rather than uniformly affecting all members.

**Figure 3:**
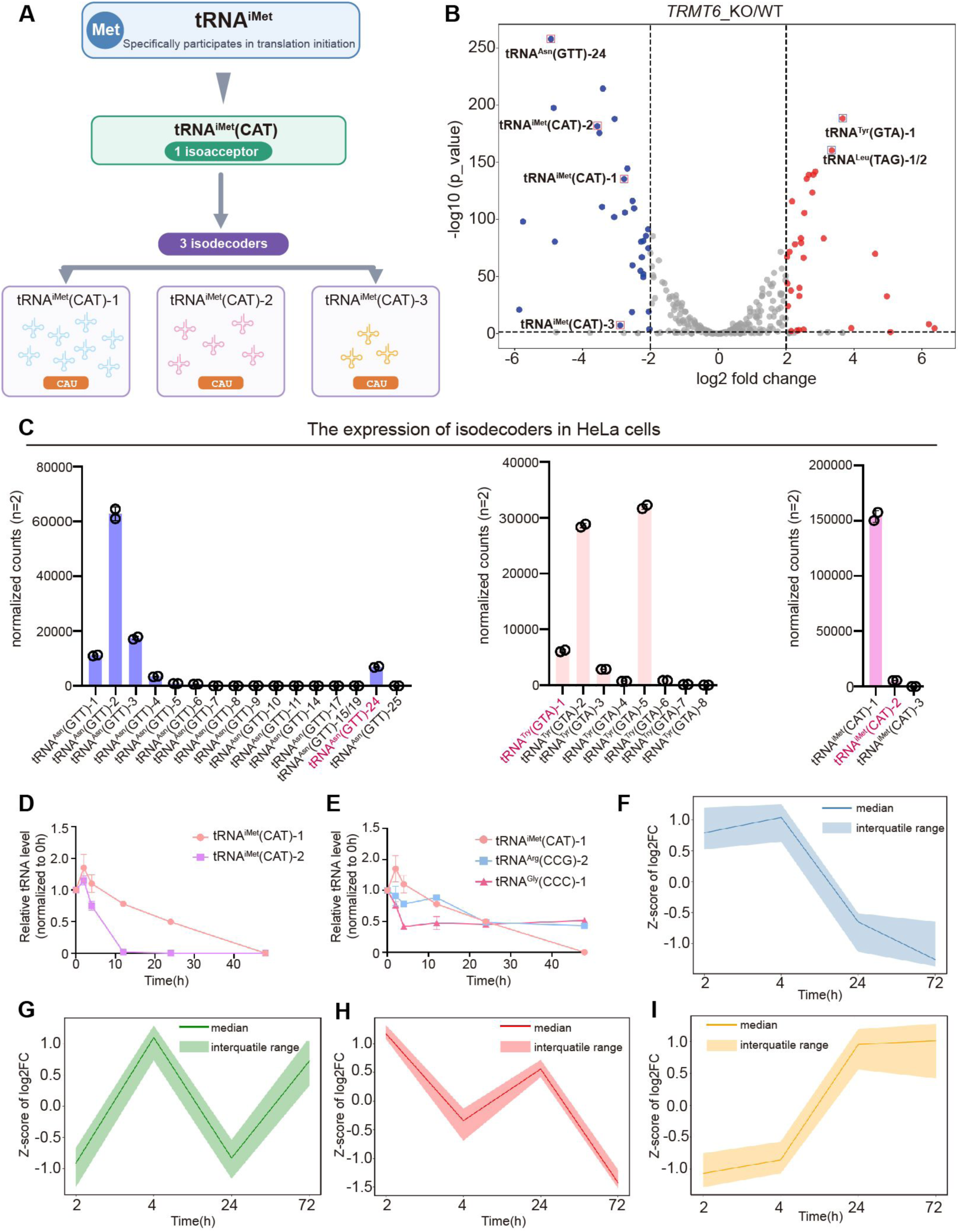
Expression, stability, and temporal response of tRNA isodecoders following TRMT6 depletion. **(A)** Schematic representation of the relationship between the isoacceptor and isodecoder classes of initiator methionine tRNA (tRNA^iMet^). Within the tRNA^iMet^ family, all members share the same methionine identity and CAT anticodon and therefore constitute one anticodon-defined isoacceptor class. Three tRNA^iMet^(CAT) sequence variants (tRNA^iMet^(CAT)-1, tRNA^iMet^(CAT)-2, and tRNA^iMet^(CAT)-3) are classified as distinct isodecoders because they differ at positions outside the anticodon. CAT denotes the DNA-encoded anticodon, corresponding to CAU in the mature tRNA. **(B)** Volcano plot showing differential tRNA isodecoder abundance between TRMT6-knockout (KO) and wild-type (WT) HeLa cells, as determined by tRNA-seq. The x-axis indicates the log2 fold change (TRMT6 KO/WT), and the y-axis indicates statistical significance (-log10(adjusted P value)). Isodecoders significantly increased or decreased following TRMT6 loss are shown in red and blue, respectively, whereas non-significant isodecoders are shown in gray. Dashed lines indicate the thresholds used to define differential abundance, and selected isodecoders are labeled. **(C)** Basal expression levels of individual tRNA isodecoders in HeLa cells, measured by tRNA-seq. Bars indicate normalized read counts, and circles represent individual biological replicates (n=2). Selected highly expressed isodecoders are highlighted. **(D)** Relative stability of tRNA^iMet^(CAT)-1 and tRNA^iMet^(CAT)-2 following transcriptional inhibition with actinomycin D. Cells were collected at the indicated time points, and isodecoder abundance was quantified by tRNA-seq and normalized to the corresponding abundance at 0 h. **(E)** Relative stability of selected highly expressed isodecoders, including tRNA^iMet^(CAT)-1, tRNA^Arg^(CCG)-2, and tRNA^Gly^(CCC)-1, following actinomycin D-mediated transcriptional inhibition. tRNA abundance at each time point was determined by tRNA-seq and expressed relative to that at 0 h. **(F-I)** Unsupervised clustering of the temporal responses of tRNA isodecoders following degron-mediated depletion of TRMT6. Based on their abundance changes at 2, 4, 24, and 72 h after TRMT6 degradation, the detected isodecoders were separated into four groups with distinct temporal patterns. **(F)** Isodecoders that remained relatively elevated at early time points but progressively decreased at 24 and 72 h. **(G)** Isodecoders showing a transient increase at 4 h, followed by a decrease at 24 h and recovery at 72 h. **(H)** Isodecoders displaying an early decrease at 4 h, a partial recovery at 24 h, and a pronounced reduction at 72 h. **(I)** Isodecoders showing low abundance at early time points followed by a sustained increase at 24 and 72 h. Solid lines represent the median Z-score of log2 fold changes within each cluster, and shaded regions indicate the interquartile range.

To determine how these isodecoder-specific changes contributed to the total abundance of each family, we next analyzed the tRNA-seq data from wild-type cells to quantify the basal abundance of each isodecoder and its relative contribution to the corresponding tRNA family (Fig. 3C; Appendix Table S2). For example, although tRNA^Asn^(GTT)-24 was significantly reduced, it represented only ∼8.8% of the total tRNA^Asn^(GTT) pool in wild-type cells (Fig. 3C; Appendix Table S2). Consequently, its reduction had only a limited effect on the overall abundance of tRNA^Asn^(GTT). Similarly, tRNA^Tyr^(GTA)-1 was expressed at a substantially lower level than the major tRNA^Tyr^(GTA) isodecoders and therefore made only a minor contribution to the total tRNA^Tyr^(GTA) pool (∼11%) (Fig. 3C; Appendix Table S2). Notably, although tRNA^Tyr^(GTA)-1 and tRNA^Tyr^(GTA)-3 were both minor members, they responded differently to *TRMT6* knockout, with a significant increase observed only for tRNA^Tyr^(GTA)-1. Thus, changes in individual isodecoders do not necessarily translate into corresponding changes at the family level, particularly when the affected isodecoder represents only a small fraction of the total pool, elucidating the global stability of the anticodon pool upon m^1^A58 loss.

To understand why tRNA^iMet^ was not buffered like other isoacceptors, we compared the basal abundance and response of its two isodecoders. tRNA^iMet^(CAT)-1 was the major contributor (∼90%), whereas tRNA^iMet^(CAT)-2 was a minor member (∼10% of the total) (Fig. 3C; Appendix Table S2). Following *TRMT6* knockout, both tRNA^iMet^(CAT)-1 and tRNA^iMet^(CAT)-2 were significantly reduced, with the minor member showing a greater proportional decline (Fig. 3B; Appendix Table S1). Thus, unlike other isoacceptor families in which stable major isodecoders remained largely unaffected, tRNA^iMet^ suffered reductions across all its detectable members and therefore lacked a buffering isodecoder to sustain its total pool.

Together, most isoacceptors are buffered by stable major isodecoders. tRNA^iMet^ lacks such a member and is therefore uniquely vulnerable to m^1^A58 loss.

### Intrinsic instability is associated with isodecoder depletion upon m^1^A58 loss

To investigate why individual isodecoders differ in their sensitivity to m^1^A58 loss, we treated cells with actinomycin D to inhibit new RNA synthesis and collected samples at multiple time points after treatment. At each time point, we used tRNA-seq to profile the entire detectable tRNA pool at isodecoder resolution, quantifying changes in the abundance of more than 350 isodecoders across seven time points relative to the 0 h time point. We found that, following actinomycin D treatment, the isodecoders most sensitive to m^1^A58 loss generally decayed faster than their corresponding major counterparts. For example, tRNA^Asn^(GTT)-24, one of the isodecoders most strongly affected by m^1^A58 loss, decayed substantially faster than the major isodecoder (tRNA^Asn^(GTT)-2) (Fig. EV3A; Appendix Table S3). Similarly, in the tRNA^Ala^(AGC) family, which contains 13 detectable isodecoders, the isodecoders most affected by m^1^A58 loss, including tRNA^Ala^(AGC)-3, -6, -15, -24, and the -14/10/12/13/16/17 group, exhibited substantially lower intrinsic stability substantially faster decay than the major isodecoder (tRNA^Ala^(AGC)-4) (Fig. EV3B, C; Appendix Table S3). These results indicate that some minor isodecoders have lower stability than their major counterparts, suggesting that the baseline tRNA stability may contribute to their differential sensitivity to m^1^A58 depletion.

Although the major tRNA^iMet^ isodecoder, tRNA^iMet^(CAT)-1, was more stable than the minor isodecoder tRNA^iMet^(CAT)-2 (Fig. 3D), we asked why it failed to buffer the m^1^A-dependent destabilization of the tRNA^iMet^ pool. Comparison with the major isodecoders of other isoacceptors showed that tRNA^iMet^(CAT)-1 was itself relatively unstable. For example, tRNA^iMet^(CAT)-1 decayed substantially faster than tRNA^Arg^(CCG)-2 and tRNA^Gly^(CCC)-1, the major isodecoders of their respective isoacceptors (Fig. 3E; Appendix Table S3). These results were further confirmed by northern blot analysis (Fig. EV2D). Thus, both tRNA^iMet^ isodecoders are sensitive to m^1^A58 loss, particularly the less stable tRNA^iMet^(CAT)-2, whereas the relatively limited stability of tRNA^iMet^(CAT)-1 restricts its capacity to buffer the loss of the tRNA^iMet^ pool.

To characterize the temporal response patterns of other isodecoders upon acute m^1^A58 loss, we analyzed the tRNA-seq data collected at 2, 4, 24, and 72 h after dTAG-13 treatment and grouped isodecoders by unsupervised clustering according to their temporal profiles. This analysis identified four groups with distinct response patterns (Fig. 3F-I). The first group remained relatively abundant at early time points but progressively decreased at 24 and 72 h (Fig. 3F). This pattern may reflect temporary buffering by the pre-existing mature tRNA pool. However, as m^1^A58 loss continues, impaired tRNA production together with ongoing turnover may eventually reduce tRNA abundance. The second group showed a transient increase at 4 h, followed by a decrease at 24 h and recovery at 72 h (Fig. 3G). This non-linear pattern suggests a transient stress response or compensatory regulation following acute TRMT6 depletion. The third group decreased at 4 h, partially recovered at 24 h, and then showed a marked reduction at 72 h (Fig. 3H). This biphasic response may result from a combination of an early direct effect, temporary compensation, and later disruption of cellular homeostasis. The fourth group showed relatively low abundance or limited changes at early time points, followed by a sustained increase at 24 and 72 h (Fig. 3I). This delayed increase is more consistent with secondary compensation or broader remodeling of the tRNA expression profile.

Together, these findings show that tRNA^iMet^ is particularly vulnerable to m^1^A58 loss because it lacks an abundant, stable, and unaffected isodecoder capable of maintaining the isoacceptor pool.

### m^1^A58 loss impairs RNase P- and RNase Z-mediated pre-tRNA^iMet^ end processing

tRNA-seq revealed that TRMT6 depletion led to accumulation of pre-tRNA^iMet^ reads with both 5’ and 3’ untrimmed ends (Fig. EV4A), suggesting that loss of m^1^A58 is associated with impaired pre-tRNA^iMet^ end maturation. To directly examine whether m^1^A58 affects pre-tRNA^iMet^ end processing, we established separate *in vitro* assays for RNase P- and RNase Z-mediated cleavage (Fig. 4A, B). For the RNase P assay, we purified endogenous human nuclear RNase P holoenzyme, a ribonucleoprotein complex comprising H1 RNA and multiple protein subunits, from Expi293F cells (Fig. 4C). For the RNase Z assay, we used recombinant human RNase Z/ELAC2 expressed in *E. coli* (Fig. 4E). Human pre-tRNA^iMet^(CAT)-1 substrates were generated by *in vitro* transcription and then methylated with SAM by the human TRMT6/TRMT61A methyltransferase complex purified from a bacterial co-expression system (Fig. EV4B). Successful m^1^A installation was confirmed by mononucleoside LC–MS/MS (Fig. EV4C). To separately assess m^1^A58 effects on RNase P- and RNase Z-mediated processing, we designed two substrates. One contained the mature tRNA^iMet^ body with both 5’ leader and 3’ trailer (as detected by tRNA-seq) and was used to monitor 5’ leader removal by RNase P. The other, a 3’-trailed pre-tRNA^iMet^ lacking the 5’ leader, served as the RNase P processing intermediate to specifically monitor 3’ trailer cleavage by RNase Z. This design decouples the two processing steps, enabling clearer assessment of m^1^A58 effects on each end.

**Figure 4:**
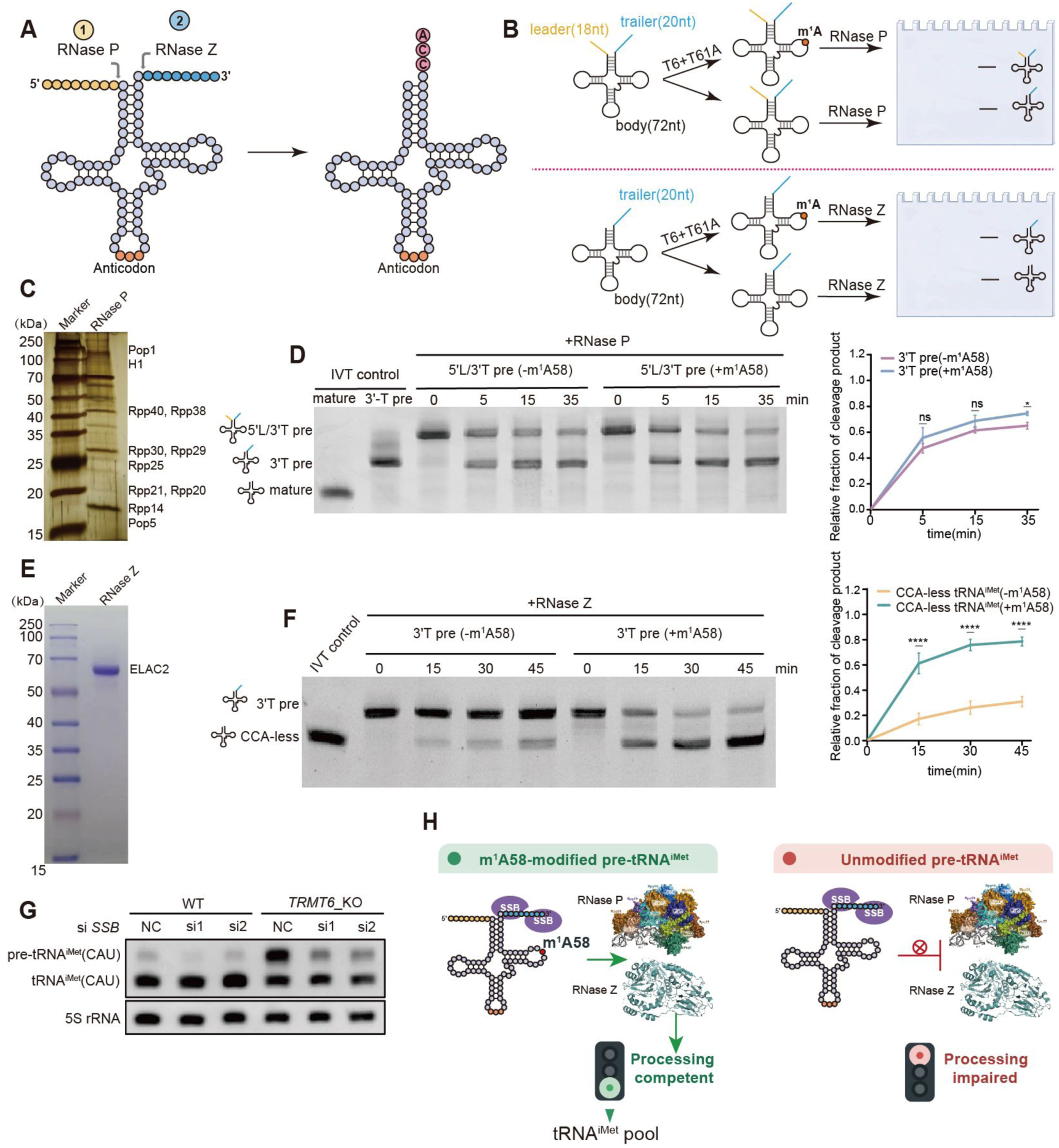
m^1^A58 loss impairs RNase P- and RNase Z-mediated pre-tRNA^iMet^ end processing. **(A, B)** Schematic showing the sequential end-processing steps of pre-tRNA maturation. RNase P removes the 5’ leader, whereas RNase Z removes the 3’ trailer to generate mature tRNA. **(C)** Silver staining of purified human RNase P holoenzyme. RNase P was purified from Expi293F cells using Rpp14 and Rpp21 as bait proteins, with co-purified subunits indicated. **(D)** Time-course *in vitro* RNase P processing assay. Full-length pre-tRNA^iMet^ (containing both 5’ leader and 3’ trailer) with or without m^1^A58 modification was incubated with purified RNase P for the indicated time points, resolved by urea-PAGE, and visualized. Mature tRNA and 3’-trailed pre-tRNA^iMet^ (lacking 5’ leader) were included as markers. Quantification of the 3’-tailed cleavage product is shown on the right, expressed as the percentage of the total input substrate remaining at each time point. **(E)** Coomassie blue staining of purified recombinant RNase Z expressed in *E. coli*. **(F)** Time-course *in vitro* RNase Z processing assay. 5’-processed pre-tRNA^iMet^ (retaining the 3’ trailer only) with or without m^1^A58 modification was incubated with purified RNase Z for the indicated time points, resolved by urea-PAGE, and visualized. Mature tRNA was included as a marker. Quantification of the mature-length cleavage product is shown on the right, expressed as the percentage of the total input substrate remaining at each time point. **(G)** Northern blot analysis of pre-tRNA^iMet^ and mature tRNA^iMet^ levels in WT and *TRMT6* KO cells with or without SSB knockdown. 5S RNA is shown as a loading control. **(H)** Proposed model. In WT cells, SSB binds and protects nascent pre-tRNA, while TRMT6/TRMT61A installs m^1^A58. m^1^A58 promotes productive handover of pre-tRNA^iMet^ to RNase P and RNase Z, enabling efficient 5’ leader and 3’ trailer removal. Upon TRMT6 loss, unmodified pre-tRNA^iMet^ remains protected by SSB but is inefficiently processed, leading to accumulation of end-unprocessed pre-tRNA^iMet^ and reduced mature tRNA^iMet^ production.

In the RNase P processing assay, we performed time-course reactions using unmodified and m^1^A58-modified 5’ leader/3’ trailer-containing pre-tRNA^iMet^. The results showed that RNase P processed the m^1^A58-modified pre-tRNA^iMet^ more efficiently (Fig. 4D). The uncleaved precursor decreased more rapidly over time, accompanied by increased production of the 3’-trailed pre-tRNA^iMet^ product after 5’ leader removal (Fig. 4D). In contrast, the pre-tRNA^iMet^ substrate lacking m^1^A58 showed a higher residual substrate fraction (Fig. 4D), indicating that loss of m^1^A58 reduces the efficiency of RNase P-mediated 5’ end processing of pre-tRNA^iMet^. Similarly, in the RNase Z processing assay, m^1^A58-modified 3’-trailed pre-tRNA^iMet^ underwent more efficient 3’-trailer cleavage and yielded higher levels of the 3’-end-processed, CCA-less tRNA^iMet^ intermediate, whereas processing of the unmodified substrate was markedly impaired (Fig. 4F). Notably, compared with RNase P-mediated 5’ leader removal, RNase Z-mediated 3’ trailer cleavage showed a stronger dependence on m^1^A58 (Fig. 4D, F). Thus, loss of m^1^A58 impairs both RNase P-mediated 5’ leader removal and RNase Z-mediated 3’ trailer cleavage, albeit with a more pronounced effect on the latter. This result is highly consistent with the tRNA-seq observation that pre-tRNA^iMet^ retaining both the 5’ leader and 3’ trailer accumulates upon m^1^A58 loss.

To understand why the unprocessed pre-tRNA^iMet^ persisted rather than being degraded, we examined the role of La/SSB, a known stabilizer of nascent RNA polymerase III transcripts (Maraia & Intine, 2001; Kufel, 2002). We depleted SSB in *TRMT6*-knockout cells and measured pre-tRNA^iMet^ levels. La/SSB depletion markedly reduced the accumulated pre-tRNA^iMet^ signal (Fig. 4G; Fig. EV5A, B). Together with the *in vitro* processing assays, these results support a model in which SSB protects pre-tRNA^iMet^ after transcription, while m^1^A58 promotes its productive entry into end processing; in the absence of m^1^A58, pre-tRNA^iMet^ remains protected but is inefficiently processed, leading to precursor accumulation and reduced mature tRNA^iMet^ production (Fig. 4H).

Together, these *in vitro* reconstitution experiments demonstrate that m^1^A58 acts early during pre-tRNA^iMet^ maturation as a processing checkpoint that promotes efficient maturation of both the 5’ and 3’ ends.

### m^1^A58 establishes a processing-competent conformation of human tRNA^iMet^

A58 is located in the T-loop of tRNA, within the D/T-loop elbow region that is critical for tRNA tertiary folding. The yeast tRNA^iMet^ structure revealed the unique initiator elbow, arrangement that differs from elongator tRNA elbow, with D/T-loop interactions involving A54, A59, A60 and A20 (Basavappa & Sigler, 1991). Structural studies of the human m^1^A58 methyltransferase revealed that it remodels the tRNA structure to access the buried A58 nucleotide (Finer-Moore *et al,* 2015). Our previous NMR studies on yeast tRNA^iMet^ showed that m^1^A58 strongly promotes proper assembly of the tRNA elbow, as indicated by the appearance of the diagnostic G18 and U55 imino signals (Yared *et al*., 2023). We therefore asked whether m^1^A58 similarly affects the conformation of human tRNA^iMet^ and thereby contributes to its recognition and processing by RNase P and RNase Z.

To examine the structural effect of m^1^A58, we compared unmodified and m^1^A58-modified human tRNA^iMet^ by NMR spectroscopy (Fig. 5A). Human tRNA^iMet^ was generated by *in vitro* transcription and methylated with purified yeast Trm6/Trm61 and SAM under conditions that ensure complete incorporation of m^1^A58. The modified RNA was refolded to obtain a homogeneous population, and 1D NMR spectra were recorded to verify that the modification was introduced quantitatively. We then recorded two-dimensional ^1^H-^15^N BEST-TROSY imino correlation spectra, which provide sensitive fingerprints of RNA base pairing, folding homogeneity, and tertiary interactions (Fig. 5A). This NMR analysis showed that the unmodified tRNA^iMet^ displayed heterogeneous and broadened imino signals, indicating conformational heterogeneity and/or exchange between alternative folding states (Fig. 5B). In contrast, m^1^A58-modified tRNA^iMet^ showed sharper and more homogeneous imino resonances, consistent with a unique and stably folded conformation (Fig. 5C). Overlay of the two spectra showed that many stem-associated imino signals were retained (Fig. 5D), suggesting that the secondary structure was largely maintained in the absence of m^1^A58 (Fig. 5E-J). Unlike yeast tRNA^iMet^ for which folding defects were only clearly identified in the elbow region (Yared *et al,* 2023), inspection of the overlay revealed additional structural defects beyond the elbow region in human tRNA^iMet^ (Fig. 5D). Signals clearly missing in the unmodified tRNA^iMet^ included the diagnostic elbow-region signals U55 and G18, as well as G15 and U8, nucleotides that form important universally conserved tertiary interactions in the tRNA core (Fig. 5B-D). Together, these missing signals indicate that both the elbow region and the tRNA core are not properly assembled in the absence of m^1^A58. Furthermore, signals from U51, G52 and G53 were also absent in the unmodified form, suggesting that the T-arm is the most affected structural element upon m^1^A58 loss (Fig. 5B-D). Thus, the structural defect caused by the lack of m^1^A58 is more pronounced than just improper elbow assembly: while most secondary structure elements are formed, the tRNA core (G15 and U8) and the T-arm (U51, G52, G53) are also compromised. However, only the m^1^A58-modified tRNA^iMet^ displayed clear imino signals corresponding to G18 and U55, and their absence in the unmodified tRNA and appearance upon modification confirm that m^1^A58 is required for proper elbow assembly in human tRNA^iMet^.

**Figure 5:**
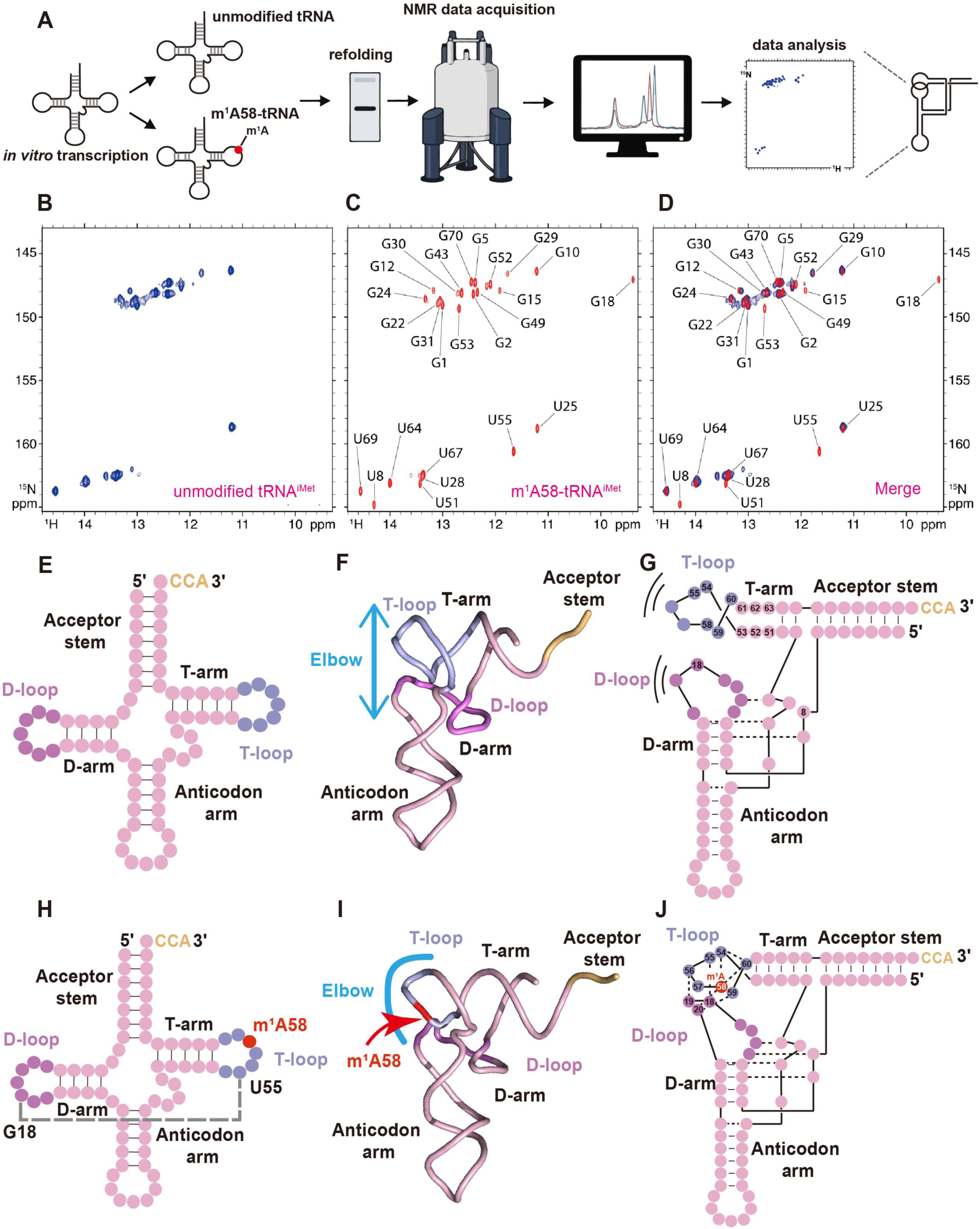
m^1^A58 establishes a processing-competent conformation of human _tRNAiMet_. **(A)** Schematic workflow for preparing unmodified and m^1^A58-modified human tRNA^iMet^ samples for NMR analysis. tRNA^iMet^ was transcribed *in vitro*, then either left unmodified or methylated at A58 using purified yeast Trm6/Trm61 and SAM. Modified tRNA was recovered by phenol-chloroform extraction and subsequently refolded to obtain a homogenous population. **(B-D)** Two-dimensional ^1^H-^15^N BEST-TROSY imino correlation spectra of human tRNA^iMet^ recorded at 38 °C. **(B)** Unmodified tRNA^iMet^ shows broadened and heterogeneous imino signals, indicative of conformational exchange and/or folding heterogeneity. **(C)** m^1^A58-modified tRNA^iMet^ displays sharper and more uniform imino resonances, consistent with a stably folded conformation. **(D)** Overlay of the unmodified (blue) and m^1^A58-modified (red) spectra shows that most imino signals from stem regions are retained. **(E-J)** Schematic models illustrating the structural consequences of m^1^A58 modification. **(E-G)** In the unmodified state, the D/T-loop elbow region is not properly assembled, precluding stable G18-U55 tertiary interactions. **(H-J)** Upon m^1^A58 methylation, the elbow architecture is stabilized, as evidenced by the formation of the G18-U55 interaction.

To assess whether this structural change affects substrate recognition by tRNA end-processing enzymes, we analyzed existing structures of the human RNase P-pre-tRNA complex and cryo-EM structures of human ELAC2 (Bhatta *et al*, 2025; Wu *et al*, 2018; Xue *et al*, 2024). In the RNase P-pre-tRNA structure, the D/T-loop elbow is recognized through the tRNA position 19/56 region, which stacks against conserved nucleotides in the H1 RNA component and helps position the acceptor stem and 5’ leader for cleavage; therefore, m^1^A58-dependent restoration of the adjacent G18-U55 elbow interaction would be expected to favor productive RNase P docking (Wu *et al*, 2018). For 3’ processing, ELAC2 structures show that the enzyme captures the T-arm/T-loop with its N-terminal flexible arm, while its C-terminal domain and C-terminal helix position the acceptor stem, A73 discriminator base, and 3’ trailer in the catalytic center, further supporting the idea that m^1^A58 stabilizes a processing-competent geometry of pre-tRNA (Bhatta *et al*, 2025; de la Sierra-Gallay *et al*, 2005; Xue *et al*, 2024). These structural observations suggest that m^1^A58 is more likely to promote processing by stabilizing the elbow architecture of pre-tRNA rather than serving as a direct recognition determinant. To understand why tRNA^iMet^ is particularly sensitive among m^1^A58-containing tRNAs, we analyzed our tRNA half-life profiling data and found that tRNA^iMet^ has one of the shortest basal half-lives across the human tRNA pool (Fig. 3D, E; Appendix Table S3). This suggests that mature tRNA^iMet^ is normally maintained under relatively high turnover. Together with previous work showing that tRNA^iMet^, but not elongator tRNAs such as tRNA^Phe^, is uniquely dependent on m^1^A58 for proper D/T-loop elbow assembly (Yared *et al*, 2023), these data suggest that tRNA^iMet^ combines an m^1^A58-dependent folding requirement with intrinsically limited basal stability. Thus, loss of m^1^A58 appears to trap tRNA^iMet^ in an elbow-incompetent state that is poorly processed, most prominently during RNase Z-mediated 3’ trailer removal. Because human tRNA^iMet^ is both unusually dependent on m^1^A58 for elbow assembly and naturally high-turnover, this folding-dependent maturation defect is selectively amplified into depletion of the mature tRNA^iMet^ pool.

Together, our data show that m^1^A58 is required for proper elbow assembly of human tRNA^iMet^. Loss of m^1^A58 preserves secondary structure but impairs processing-competent folding, especially RNase Z-mediated 3’ cleavage. Because tRNA^iMet^ relies heavily on m^1^A58 for folding and has a short half-life, this defect depletes the mature tRNA pool.

### m^1^A58-dependent tRNA^iMet^ maturation supports translation initiation and shapes temporal gene-expression responses

During translation initiation, eIF2-GTP delivers tRNA^iMet^ to the 40S ribosomal subunit to form the 43S pre-initiation complex (PIC) (Hinnebusch, 2014; Sonenberg & Hinnebusch, 2009). Our preceding results showed that loss of m^1^A58 modification on tRNA^iMet^ impairs its precursor processing and reduces the abundance of mature tRNA^iMet^. To determine whether this maturation defect compromises protein synthesis, we measured puromycin incorporation in *TRMT6*-knockout cells. Puromycin incorporation was markedly reduced following TRMT6 loss (Fig. 6A), indicating decreased nascent protein synthesis. Supplementation with m^1^A58-modified tRNA^iMet^ substantially restored puromycin incorporation (Fig. 6B, C), demonstrating that the translational defect is primarily attributable to an insufficient pool of functional tRNA^iMet^ rather than nonspecific cellular damage.

**Figure 6:**
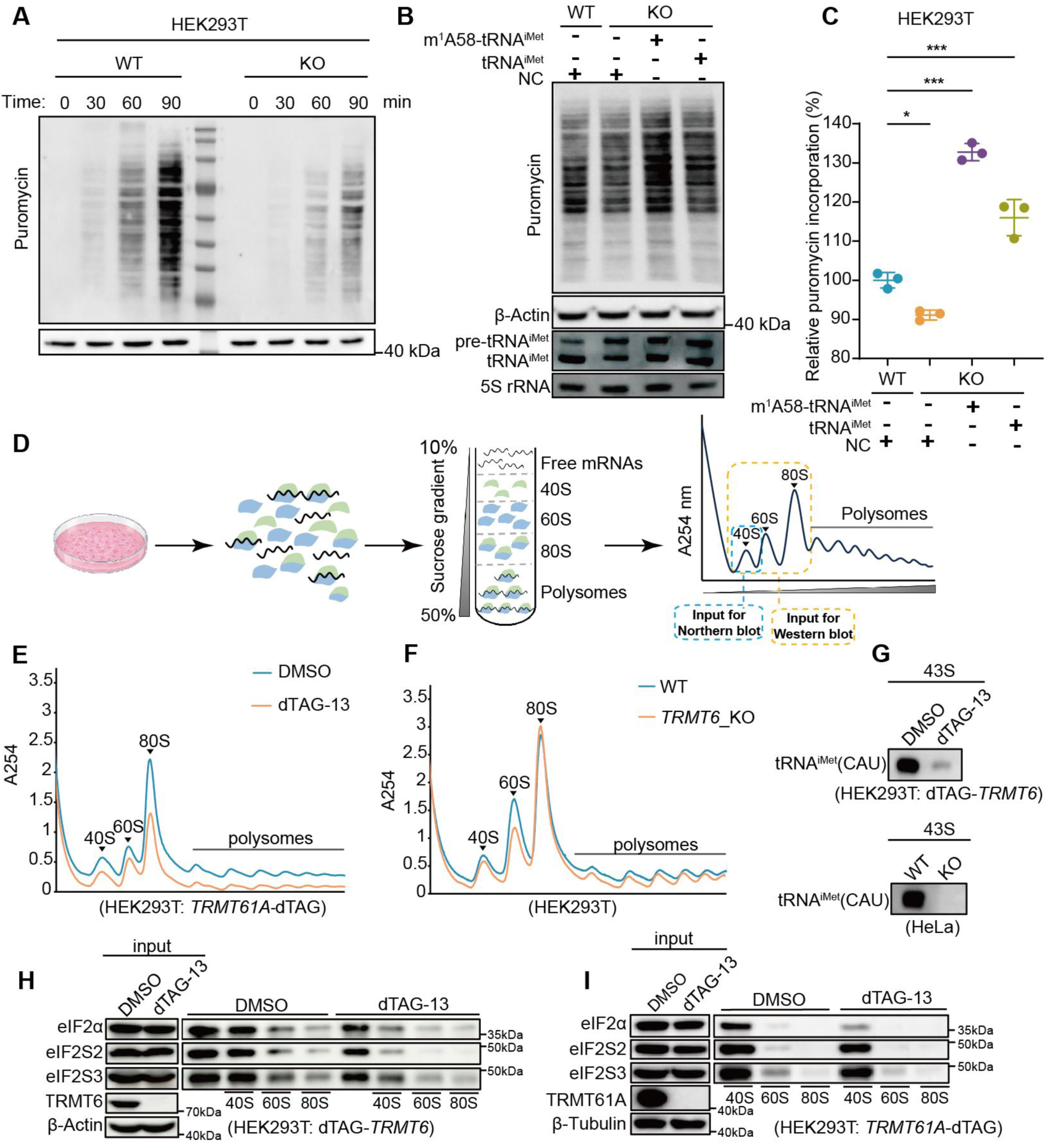
m^1^A58-dependent maturation of tRNA^iMet^ supports global protein synthesis, polysome formation, and 43S pre-initiation complex assembly. **(A)** Puromycin-incorporation assay measuring nascent protein synthesis in wild-type (WT) and *TRMT6*-knockout (KO) HEK293T cells. Cells were pulse-labeled with puromycin for the indicated times, and incorporated puromycin was detected by immunoblotting with an anti-puromycin antibody. **(B)** Rescue of protein synthesis in *TRMT6*-KO cells by exogenous tRNA^iMet^. *TRMT6*-KO cells were transfected with *in vitro*-transcribed unmodified tRNA^iMet^(CAU) or m^1^A58-modified tRNA^iMet^(CAU), followed by puromycin labeling and anti-puromycin immunoblotting. Mock-transfected *TRMT6*-KO cells served as the control. **(C)** Quantification of the puromycin-incorporation signals shown in **(B)**. Puromycin signals were normalized to the WT. Statistical significance was determined using t-test. *P < 0.05; **P < 0.001. **(D)** Schematic of polysome profiling used to assess ribosome loading onto mRNAs. Cytoplasmic extracts were separated by sucrose-density-gradient ultracentrifugation, and absorbance at 254 nm was continuously monitored to resolve free ribosomal subunits, monosomes, and polysomes. **(E)** Representative polysome profiles of control and dTAG-treated cells following acute degradation of TRMT61A. Peaks corresponding to the 40S and 60S ribosomal subunits, 80S monosomes, and polysomes are indicated. **(F)** Representative polysome profiles of WT and TRMT6_KO cells. **(G)** Analysis of tRNA^iMet^(CAU) associated with the 43S pre-initiation complex. Cells were mildly crosslinked with formaldehyde to stabilize transient initiation complexes, and 43S-containing fractions were isolated by ultracentrifugation. tRNA^iMet^(CAU) in the isolated fractions was examined after dTAG-induced TRMT6 degradation in HEK293T cells and in *TRMT6*-KO HeLa cells. The corresponding control cells were processed in parallel. **(H, I)** Immunoblot analysis of translation-initiation factors associated with isolated 43S fractions following dTAG-induced degradation of TRMT6 **(H)** or TRMT61A **(I)**. Cells were mildly crosslinked with formaldehyde before 43S complexes were isolated by ultracentrifugation. The abundance of the indicated 43S-associated proteins, including eIF2α, eIF2S2, and eIF2S3, was assessed by Western blotting.

To determine whether impaired protein synthesis results from defective translation initiation, we performed polysome profiling in *TRMT6*-knockout cells and in cells subjected to 48 h of dTAG-13-induced degradation of TRMT6 or TRMT61A (Fig. 6D). All three perturbations reduced polysome-associated fractions, consistent with decreased productive ribosome loading onto mRNAs (Fig. 6E, F). To examine 43S PIC assembly more directly, we stabilized transient initiation complexes by mild formaldehyde crosslinking and isolated the 43S fraction. We first confirmed that mature tRNA^iMet^ was enriched in this fraction, and its level was markedly reduced upon TRMT6 or TRMT61A degradation (Fig. 6G). Consistently, Western blot analysis showed that TRMT6 or TRMT61A degradation markedly reduced the recovery of eIF2α, eIF2β, and eIF2γ in this fraction without a corresponding reduction in their total abundance (Fig. 6H, I). Thus, TRMT6 or TRMT61A loss specifically impairs eIF2 association with the 43S complex, supporting a requirement for m^1^A58-dependent tRNA^iMet^ maturation in efficient PIC assembly.

Having established that loss of TRMT6 impairs tRNA^iMet^ maturation and compromises protein synthesis, we next asked how cells respond transcriptionally to sustained translation-initiation stress. We performed time-course RNA-seq at 2, 6, 12, and 24 h after dTAG-13-induced TRMT6 degradation. Clustering based on normalized transcript counts identified two broad temporal expression patterns (Fig. 7A, B). Count-C1 genes increased from 2 to 12 h and declined by 24 h (Fig. 7A), whereas Count-C2 genes decreased through 12 h and partially recovered at 24 h (Fig. 7B). Clustering by fold change relative to 0 h further resolved four response patterns (Fig. 7C-F). FC-C1 genes increased between 2 and 12 h before declining at 24 h and were enriched in the unfolded protein response (Fig. 7C), post-translational protein folding, and chaperone-dependent refolding (Fig. 7G). FC-C2 genes decreased at 6-12 h and rebounded by 24 h and were associated predominantly with amino acid transport and cytokine-regulatory processes (Fig. 7D, G). FC-C3 genes declined progressively through 12 h before partially recovering (Fig. 7E), whereas FC-C4 genes showed an early decrease followed by sustained recovery through 24 h (Fig. 7F). These latter clusters were enriched in processes related to cell migration, signaling, transcriptional regulation, and development (Fig. 7G). Thus, TRMT6 degradation induces a temporally structured transcriptional response characterized by dynamic regulation of proteostasis, transport, signaling and regulatory pathways rather than a uniform change in gene expression. Consistent with the acute degron response, Reactome pathway analysis of differentially expressed genes in *TRMT6*-knockout cells highlighted proteostasis-related pathways, including the unfolded protein response and IRE1α- and HSF1/HSP90-associated programs (Fig. 7G). Thus, both acute and sustained TRMT6 loss elicits a shared proteostasis-stress response.

**Figure 7:**
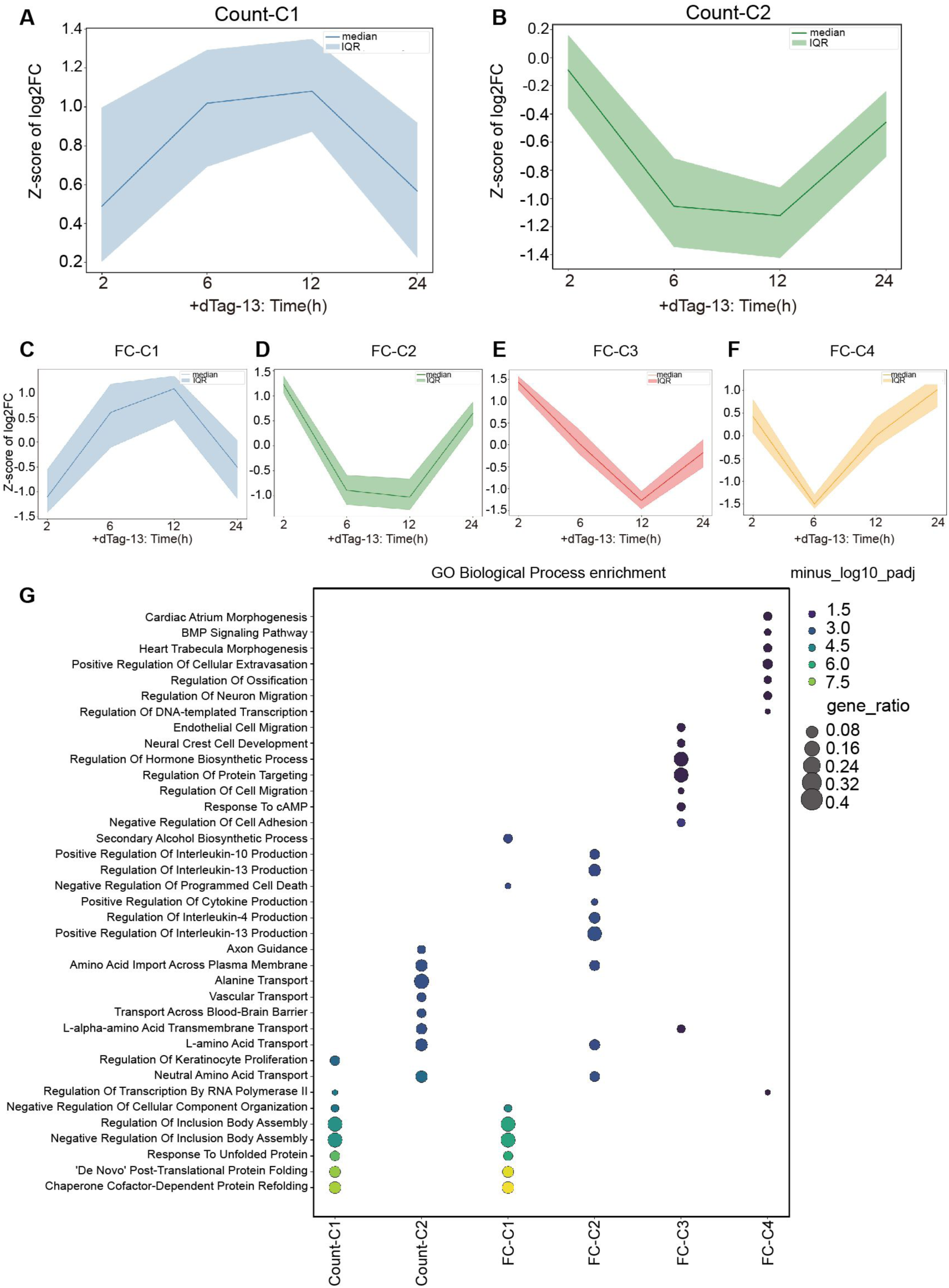
Temporal gene-expression patterns following dTAG-13-induced degradation of TRMT6. **(A, B)** Temporal expression profiles of genes grouped into two clusters (Count-C1 and Count-C2) based on normalized RNA-seq counts at the indicated time points after dTAG-13 treatment. **(C-F)** Genes grouped into four clusters (FC-C1-FC-C4) according to their temporal fold-change (FC) profiles relative to 0 h, illustrating distinct time points at which gene-expression trends change. In **(A-F)**, normalized count-derived expression values or log2-transformed FC values were standardized as Z-scores. Solid lines indicate the median, and shaded areas indicate the interquartile range (IQR) of genes within each cluster. **(G)** Gene Ontology (GO) Biological Process enrichment analysis of genes in each count- or FC-based cluster. Dot size represents the gene ratio, and dot color indicates enrichment significance, shown as − log10 of the adjusted P value (− log10 Padj). h, hours; FC, fold change; IQR, interquartile range; GO, Gene Ontology; Padj, multiple-testing-adjusted P value.

Collectively, these findings establish m^1^A58-dependent tRNA^iMet^ maturation as a critical link between tRNA biogenesis and translation-initiation capacity, while revealing a temporally coordinated gene-expression response to acute TRMT6 loss.

## Discussion

tRNA modifications are viewed as chemical marks that regulate the stability, decoding capacity, or translational efficiency of mature tRNAs (Orellana *et al*, 2021; Sakai *et al*, 2019; Suzuki, 2021). We show that m^1^A58 promotes human initiator tRNA to adopt a conformation competent for end processing, thereby linking tRNA folding and maturation to translation initiation capacity. Although m^1^A58 is widely distributed among human cytoplasmic tRNAs, acute or stable loss of TRMT6 caused only modest changes in the overall tRNA pool. This apparent robustness can be explained, at least in part, at the isodecoder level: for most isoacceptors, the isodecoders affected by TRMT6 loss are low-abundance members, whereas abundant and stable major isodecoders remain largely unchanged and buffer the total pool. This is reminiscent of a similar buffering mechanism observed during hiPSC differentiation, where stable major isodecoders maintain anticodon pool homeostasis despite extensive reprogramming of the tRNA transcript repertoire (Gao *et al*, 2024). By contrast, tRNA^iMet^ lacks a stable and unaffected isodecoder. Its major isodecoder is not intrinsically stable and is sensitive to TRMT6 loss; the minor isodecoder is even more short-lived, and the third is expressed near the detection limit. This absence of a truly stable isodecoder makes the initiator-tRNA pool particularly vulnerable to disruption of m^1^A58 deposition.

This vulnerability is likely rooted in a maturation defect: if m^1^A58 is required for efficient end processing, its loss would impair production of functional tRNA^iMet^. Consistent with this view, biochemical reconstitution showed that m^1^A58 enhanced both RNase P-mediated 5’-leader removal and RNase Z-mediated 3’-trailer cleavage. The effect was particularly pronounced for RNase Z, suggesting that the geometry required for productive 3’-end processing may be especially sensitive to the m^1^A58-dependent structural state of the tRNA elbow. These findings establish a direct role for m^1^A58 in promoting pre-tRNA^iMet^ end maturation. Our NMR analyses provide a structural explanation for this processing defect. Loss of m^1^A58 did not abolish the overall secondary structure of human tRNA^iMet^, as many stem-associated imino resonances were retained. Instead, the unmodified tRNA exhibited broader and more heterogeneous signals, whereas m^1^A58 installation produced a more homogeneous spectrum and restored the diagnostic G18 and U55 imino signals associated with proper D/T-loop elbow assembly. Thus, m^1^A58 appears to shift tRNA^iMet^ from a more dynamic ensemble toward a more stably folded, processing-competent state. Because both RNase P and RNase Z engage structural features surrounding the T-arm and tRNA elbow, this conformational stabilization provides a plausible mechanism by which m^1^A58 promotes productive substrate recognition and cleavage. Rather than functioning as a direct sequence-specific recognition determinant, m^1^A58 likely facilitates processing by establishing the global geometry required for enzyme engagement.

Our transcription-shutoff experiments showed that the major tRNA^iMet^ isodecoder is less stable than representative major isodecoders from other isoacceptors. Consequently, even a moderate reduction in productive tRNA^iMet^ maturation may rapidly deplete the functional mature pool because existing molecules are continuously turned over. However, the pathway responsible for eliminating unprocessed or hypomodified tRNA^iMet^ has not been defined here. It is worth noting that Xrn1 and Xrn2 have been shown to mediate tRNA^iMet^ degradation in heat-stressed HeLa cells, with Xrn1 acting in the cytoplasm and Xrn2 in the nucleus (Watanabe *et al*, 2013). Identifying the surveillance factors that recognize these structurally altered tRNA species will therefore be an important direction for future work.

The selective depletion of mature tRNA^iMet^ provides a direct connection between tRNA maturation and translation initiation. Initiator Met-tRNA^iMet^ is delivered to the 40S ribosomal subunit by eIF2-GTP and is therefore an essential component of the 43S pre-initiation complex (Shin *et al*, 2011). Consistent with reduced availability of functional tRNA^iMet^, TRMT6 or TRMT61A depletion reduced polysome abundance, decreased tRNA^iMet^ recovery in the 43S fraction, and impaired the association of eIF2 subunits with this fraction without reducing their total cellular abundance. Moreover, supplementation with m^1^A58-modified tRNA^iMet^ substantially restored puromycin incorporation. These results support the conclusion that defective tRNA^iMet^ maturation is a major contributor to the translational phenotype caused by loss of the TRMT6/TRMT61A complex. Given that overexpression of tRNA^iMet^ has been shown to promote cell proliferation and transformation in breast epithelial cells (Pavon-Eternod *et al*, 2013), our findings raise the possibility that the m^1^A58-dependent maturation of tRNA^iMet^ may represent a previously unappreciated node for therapeutic intervention in cancers that depend on this pathway.

More broadly, TRMT6/TRMT61A has emerging functions beyond tRNA modification, including the regulation of tRNA-derived fragments, and a non-canonical methyltransferase-independent role in hematopoietic stem-cell homeostasis (Su *et al*, 2022; He *et al*, 2024). Its activity also extends to tRNA-like and non-canonical RNA substrates, such as the MALAT1-derived mascRNA and CAG-repeat RNAs (Skeparnias *et al*, 2024; Sun *et al*, 2023), further highlighting the diverse biological roles of this modification machinery.

## Methods

### Cell lines and culture

HEK293T and HeLa cells were cultured in DMEM with 10% fetal bovine serum and 1% penicillin/streptomycin at 37 °C and 5% CO_2_.

### Generation of TRMT6 knockout cell lines and Western blotting verification

TRMT6 knockout cell lines were established in HEK293T and HeLa cells using sgRNAs targeting the exonic region of human TRMT6. The sgRNA sequences were designed via the CRISPR online tool (http://crispr.mit.edu) and inserted into the pX330-mCherry vector. The two sgRNA sequences used were as follows:

sgRNA-1: 5’-CACCGCGAAGTCGCCGTCGCGGATG-3’

sgRNA-2: 5’-CACCGAGTCGCCGTCGCGGATGCGG-3’

Annealed sense and antisense oligonucleotides were ligated into the pX330-mCherry vector, which expresses both Cas9 nuclease and mCherry. Recombinant plasmids were purified by endotoxin-free midiprep and transfected into host cells. At 36 h post-transfection, mCherry-positive cells were sorted by flow cytometry, and single positive cells were seeded into 96-well plates for monoclonal expansion.

After sufficient proliferation, genomic DNA was extracted from individual clones and subjected to PCR amplification using locus-specific primers. PCR products were preliminarily examined by GelRed-stained agarose gel electrophoresis. Sanger sequencing was further performed to verify genome-editing efficiency and mutation patterns. Western blotting was performed to confirm the loss of endogenous protein expression using anti-TRMT6 (Proteintech, 16727-1-AP, 1:2000) and anti-TRMT61A (Abcam, A25353, 1:2000) antibodies.

### Plasmid construction

The previously reported CRISPR-Cas9 backbone plasmid pX330-mcherry and pTrc99b donor plasmid were used as the fundamental vectors in this study.

The custom-constructed plasmids included a pX330-sgRNA vector targeting the N-terminus of human TRMT6 (sgRNA sequence: 5’-GACCGGCTGAGCGTCATGGA-3’), and a matching pTrc99b donor plasmid for knock-in at the TRMT6 start codon locus, which contains upstream 672 bp and downstream 959 bp homologous arms.

In addition, a pX330-sgRNA vector targeting the C-terminus of human TRMT61A (sgRNA sequence: 5’-CACCAAGACCCCAGGCTAGG-3’) and a corresponding pTrc99b knock-in donor plasmid were generated. This donor plasmid flanks the TRMT61A stop codon region with 1997 bp upstream and 581 bp downstream homologous arms.

### dTAG-mediated protein degradation

FKBP12 PROTAC dTAG-13 (MedChemexpress, HY-114421) and its negative control dTAG-13-NEG (MedChemexpress, HY-114421) were dissolved in dimethyl sulfoxide (DMSO) to prepare 1 mM stock solutions. Cells were treated with dTAG-13 at a final concentration of 0.4 μM for the indicated durations to induce targeted protein degradation, with DMSO-treated cells serving as the vehicle control. For long-term treatment exceeding 3 days, fresh complete medium containing the corresponding compound was replenished to maintain stable drug concentration.

### Northern blotting

Northern blotting was performed according to previously described protocols. Briefly, 1-2 μg of total RNA was separated on 12% urea-polyacrylamide gels containing 7 M urea and 1× TBE buffer. Fractionated RNA was transferred to positively charged Roche nylon membranes (Merck, 11417240001), The membrane was crosslinked at 254 nm, with an energy of 150 mJ/cm^2^ using UV Stratalinker 1800. Membranes were then hybridized overnight at 55 °C with 3’-DIG-labeled DNA probes at a final amount of 50 pmol per membrane. All custom DNA probes were synthesized by Sangon Biotech (Shanghai, China). DIG-labeled probes were designed to detect tRNA, with 5S rRNA serving as the internal loading control. Probe sequences were designed based on tRNA gene sequences retrieved from the GtRNAdb 2.0 database (Chan & Lowe, 2016)

For DIG signal development, membranes were incubated with anti-DIG-alkaline phosphatase-conjugated antibody (Merck, 11093274910), and chemiluminescent signals were generated using CDP-Star substrate (Merck, 12041677001). Blot signals were captured using the Amersham™ ImageQuant™ imaging system (Cytiva). Band intensities were quantified with ImageJ software. Unless otherwise specified, the signal intensities of target tRNAs were normalized to the 5S rRNA signal. All DNA probe sequences are listed in Appendix Table S4.

### TRMT6-TRMT61A expression and purification

The human *TRMT6* (NM_015939.5) was inserted into the pETDuet-1 vector with an N-terminal 6× His-tag via restriction enzyme digestion, and the *TRMT61A* (NM_152307.3) was subsequently inserted into the multiple cloning site (MCS) of the same vector by homologous recombination. The resulting recombinant plasmid pETDuet-1-TRMT6-TRMT61A was transformed into *Escherichia coli* Rosetta competent cells. Protein expression was induced overnight at 18 °C with 0.2 mM IPTG.

Bacterial cells were harvested and resuspended in lysis buffer containing 20 mM Tris-HCl (pH 7.5), 500 mM NaCl, 5 mM MgCl_2_, 5 mM DTT, 10 mM imidazole, and 10% glycerol. Cells were ultrasonicated on ice in the presence of 1 mM PMSF. The lysate was centrifuged at 16,000 g for 30 min at 4 °C, and the supernatant was incubated with Ni NTA Beads (Smart, SA005025).

The beads were sequentially washed with lysis buffer, high-salt buffer containing 1 M NaCl, and buffer supplemented with 15 mM and 25 mM imidazole. The 6X His-tagged TRMT6–TRMT61A complex was finally eluted using 250 mM imidazole. Eluted proteins were dialyzed against dialysis buffer (20 mM Tris-HCl, pH 7.5, 200 mM NaCl, 5 mM MgCl_2_, 2 mM DTT), concentrated, and supplemented with 50% glycerol for long-term storage at -20 °C.

### Preparation of tRNA and pre-tRNA substrates

The tRNA and pre-tRNA substrates used in this study, including tRNA^iMet^(CAU) and pre-tRNA^iMet^(CAU)-1-8, were generated by *in vitro* transcription (Li *et al*, 2021). Target sequences were cloned into the pTrc99b vector using BamHI and EcoRI restriction enzymes, and *in vitro* transcription was performed using T7 RNA polymerase.

Transcription products were purified by urea-denaturing polyacrylamide gel electrophoresis. RNAs were eluted with 0.5 M sodium acetate (pH 5.2) and precipitated with ethanol overnight at -20 °C. Purified RNAs were resuspended in 5 mM MgCl₂, annealed at 85 °C for 5 min, and cooled naturally to room temperature to facilitate correct tRNA refolding.

### *In vitro* tRNA methylation assay

*In vitro* methylation assays were performed to prepare tRNA/pre-tRNA substrates with (+m^1^A58) or without (-m^1^A58) m^1^A58 modification, using *in vitro* transcribed and purified tRNA/pre-tRNA as substrates.

For +m^1^A58 sample preparation, 8 μg of tRNA/pre-tRNA was incubated with 2 μM TRMT6-TRMT61A complex in a 200 μL reaction system (50 mM Tris-HCl, pH 7.5, 200 mM NaCl, 5 mM MgCl_2_, 100 μg/mL BSA, 5 mM DTT, 200 μM SAM) at 37 °C for 1 h.

After incubation, RNA was purified by phenol-chloroform extraction and ethanol precipitation. The obtained RNA pellets were washed with 75% ethanol, air-dried and resuspended in RNase-free water for subsequent experiments.

For the -m^1^A58 control group, equal amounts of batch-matched tRNA/pre-tRNA were incubated in an identical buffer system under the same conditions without TRMT6/TRMT61A complex. Identical RNA purification procedures were performed to eliminate experimental interference.

The m^1^A58 modification levels of all prepared RNA samples were quantified via LC-MS/MS analysis.

### RNase P cleavage assay

The pLVX-RPP21-Twin-Strep-IRES-EGFP and pLVX-RPP14-3×Flag-IRES-mCherry plasmids used in this study were kindly provided by Professor Ming Lei’s group. Expi293F cells were transfected with the above plasmids, cultured in Union-293 medium (Union-Biotech, UP0050), and sorted by flow cytometry. Human RNase P protein was purified according to previously reported protocols.

For RNase P cleavage reactions, 150 ng of folded pre-tRNA with or without m1A modification was incubated with 200 ng of RNase P protein in a 10 μL reaction buffer containing 25 mM Tris-HCl (pH 7.5), 100 mM NaCl, and 10 mM MgCl_2_. Reactions were performed for 0, 5, 15, and 35 min respectively and terminated by adding 10 μL of 2× RNA loading buffer followed by heating at 85 °C for 5 min.

Samples were separated by 10% TBE-Urea gel electrophoresis. Gels were stained with SYBR Gold nucleic acid dye for 5 min and imaged at 254 nm excitation using a Tanon 2500/2500R fully automatic gel imaging system. Band intensities were quantified using ImageJ software for subsequent statistical analysis. All experiments were performed with three independent biological replicates.

### RNase Z cleavage assay

A truncated human ELAC2 protein comprising amino acid residues 50-826 was produced using a baculovirus-insect cell expression system. The coding sequence corresponding to amino acid residues 50-826 of human *ELAC2* (NM_018127.7) was cloned into a modified pFastBac1 vector to generate a recombinant protein bearing an N-terminal His tag. The verified recombinant plasmid was transformed into DH10Bac competent cells. Positive clones were screened by blue-white selection, and recombinant Bacmid DNA was extracted. ELAC2 protein expression and purification were performed according to previously published protocols.

For RNase Z cleavage assays, 150 ng of folded mature tRNA was incubated with 0.1 μM ELAC2 protein in a 10 μL reaction buffer containing 25 mM Tris-HCl (pH 7.5), 100 mM NaCl, and 10 mM MgCl_2_. Reactions were terminated at 0, 15, 30, and 45 min by adding 10 μL of 2× RNA loading buffer followed by heating at 85 °C for 5 min. Samples were separated by 10% TBE-Urea gel electrophoresis.

Gels were stained with SYBR Gold nucleic acid dye for 5 min and imaged at 254 nm ultraviolet excitation using a Tanon 2500/2500R automatic gel imaging system. Band intensities were quantified via ImageJ software for subsequent statistical analysis. All experiments were performed with three independent biological replicates.

### Protein extraction and Western blotting

Western blotting was performed according to a previously described protocol. Whole-cell proteins were extracted from cultured cells or tissue samples using RIPA lysis buffer (Merck, 20-188) supplemented with 1% Protease Inhibitor Cocktail (MedChemexpress, HY-K0012) to prevent protein degradation. Equal amounts of protein samples were separated by 10% SDS-PAGE gel electrophoresis. Target proteins were immunodetected with specific primary antibodies, the detailed information of which is listed in Appendix Table S4. For secondary antibody incubation, HRP-conjugated goat anti-mouse IgG (H+L) (Yeasen, 33201ES60, 1:5000 dilution) and HRP-conjugated goat anti-rabbit IgG (H+L) (Yeasen, 33101ES60, 1:5000 dilution) were applied for corresponding target detection.

Blots were visualized using the Omni-ECL Femto Light Chemiluminescence Kit (EpiZyme, SQ202L), and chemiluminescent signals were captured with the Amersham ImageQuant imaging system (Cytiva). Band intensities were quantified using ImageJ software. The relative expression levels of target proteins were normalized to β-actin as the internal reference.

### Puromycin incorporation

Cells were subjected to puromycin pulse labeling over a time course ranging from 0 mim to 60 mim or 90 min. Cells were subsequently lysed, and whole-cell protein extracts were analyzed via Western blotting using an anti-puromycin antibody to assess global translational activity. β-actin was detected in parallel as the loading control.

### Polysome profiling and pre-initiation complex (PIC) analysis

For polysome profiling, cells were pre-treated with cycloheximide prior to lysis. Whole-cell lysates were loaded onto 10-50% linear sucrose gradients and separated by ultracentrifugation, with continuous absorbance monitoring at 254 nm (A254) to generate ribosome distribution profiles.

For PIC analysis, cells were crosslinked on ice with 0.04% formaldehyde for 5 min without cycloheximide pre-treatment. Cell lysates were fractionated on identical 10-50% sucrose gradients. Individual gradient fractions were subsequently subjected to Western blotting to probe translational initiation factors and Northern blotting to detect tRNA^iMet.^

### tRNA^iMet^ supplementation

*In vitro*-transcribed tRNA^iMet^ with or without m^1^A58 was transfected into *TRMT6* KO HEK293T cells with Lipofectamine, and puromycin incorporation was measured to assess translational rescue.

### Preparation of unmodified and m^1^A58-modified human tRNA^iMet^ for NMR

Unmodified human tRNA^iMet^ was prepared by standard *in vitro* transcription following previously published procedures, either with unlabelled NTPs or ^15^N-labelled Us and Gs (Gato *et al*, 2021; Yared *et al,* 2024). We replaced the first Watson Crick base pair A1-U72 of tRNA^iMet^ with a G1-C72 base pair in order to improve *in vitro* transcription efficiency. Briefly, tRNA^iMet^ was *in vitro* transcribed with T7 polymerase and purified by anion exchange chromatography. To prepare the single modified m^1^A58-tRNA^iMet^, 46 nmol of refolded tRNA^iMet^ was incubated overnight at 30 °C with 4.6 nmol of purified yeast Trm6/Trm61 and 500 nmol of S-adenosyl-L-methionine (SAM). The modification reaction was performed in the following maturation buffer (MB): 100 mM NaH_2_PO_4_/K_2_HPO_4_ pH 7.0, 5 mM NH_4_Cl, 2 mM DTT and 0.1 mM EDTA. Then, the tRNA product was extracted using standard phenol extraction procedures, dialyzed extensively against Na-phosphate pH 6.5 1 mM, refolded by heating at 95 °C for 5 min and cooled down slowly at room temperature. Buffer was added to place the tRNA^iMet^ in the NMR buffer: 10 mM Na-phosphate pH 6.5, 10 mM MgCl_2_, and the sample was concentrated using Amicon 10,000 MWCO (Millipore).

### NMR spectroscopy

All NMR spectra of human tRNA^iMet^ were measured at 38 °C on a Bruker AVIII-HD 700 MHz spectrometer equipped with TCI 5-mm cryoprobe with 5-mm Shigemi tubes in the NMR buffer (10 mM Na-phosphate pH 6.5, 10 mM MgCl_2_) supplemented with 5% (v/v) D_2_O. To verify that the desired modification (m^1^A58) was incorporated quantitatively in human tRNA^iMet^, 1D jump-and-return-echo NMR spectra (Plateau *et al*, 1982; Sklenar *et al*, 1987) of the modified and unmodified tRNAs were measured and compared to previously characterized samples (Yared *et al*, 2023). 2D (^1^H,^15^N)-BEST-TROSY spectra (Farjon *et al*, 2009) of unmodified tRNA^iMet^ and m^1^A58-tRNA^iMet^ were measured to evaluate the effect of m^1^A58 on the structural properties of human tRNA^iMet^. Imino resonances of the m^1^A58-tRNA^iMet^ were assigned using 2D jump-and-return-echo (^1^H,^1^H)-NOESY (Plateau *et al*, 1982; Sklenar *et al*, 1987) and 2D (^1^H,^15^N)-BEST-TROSY experiments. The data were processed using TOPSPIN 3.6 (Bruker) and analysed with NMRFAM-SPARKY (Lee *et al*, 2015).

### mRNA seq and tRNA seq analysis

#### Preprocessing and alignment of tRNA-seq reads

Sequencing reads were processed using Trim Galore (v0.6.10), incorporating Cutadapt (v4.4) and FastQC (v0.12.1), for adapter and poly(A)-tail removal and quality control. Reads were filtered using a minimum Phred score of 20 and a minimum length of 20 bp. tRNA reads were mapped to the human reference transcriptome using mim-tRNAseq (v1.3.11), with an initial cluster identity threshold of 0.95.

#### tRNA abundance and modification analysis

tRNA abundance differentiational analysis use pyDeseq2 (0.4.9) for isodecoder count and isoacceptor count files. Modification signatures, read stops and readthrough profiles were extracted from mim-tRNAseq output.

#### mRNA seq processing

mRNA seq were processed using Trim Galore (v0.6.10), incorporating Cutadapt (v4.4) and FastQC (v0.12.1), for adapter removal and quality control. Then samples were mapped to GRCh38 genome with STAR (2.7.11a), features were count with featureCounts (v2.0.6) with -t exon -g gene_id. mRNA differentiational analysis use pyDeseq2 (v0.4.9). Functional enrichment analysis was performed using Enrichr through the gseapy package. Differentially expressed genes were analyzed against the MSigDB Hallmark 2020 and Gene Ontology Biological Process 2023 gene-set collections using the human gene annotation.

